# On estimating evolutionary probabilities of population variants

**DOI:** 10.1101/475475

**Authors:** Ravi Patel, Sudhir Kumar

**Affiliations:** Institute for Genomics and Evolutionary Medicine, Temple University; Department of Biology, Temple University

**Keywords:** generalized method, evolutionary probability, forbidden alleles, potential adaptation

## Abstract

**Background:** The evolutionary probability (EP) of an allele in a DNA or protein sequence predicts evolutionarily permissible (ePerm; EP ≥ 0.05) and forbidden (eForb; EP < 0.05) variants. EP of an allele represents an independent evolutionary expectation of observing an allele in a population based solely on the long-term substitution patterns captured in a multiple sequence alignment. In the neutral theory, EP and population frequencies can be compared to identify neutral and non-neutral alleles. This approach has been used to discover candidate adaptive polymorphisms in humans, which are eForbs segregating with high frequencies. The original method to compute EP requires the evolutionary relationships and divergence times of species in the sequence alignment (a timetree), which are not known with certainty for most datasets. This requirement impedes a general use of the original EP formulation. Here, we present an approach in which the phylogeny and times are inferred from the sequence alignment itself prior to the EP calculation. We evaluate if the modified EP approach produces results that are similar to those from the original method.

**Results:** We compared EP estimates from the original and the modified approaches by using more than 18,000 protein sequence alignments containing orthologous sequences from 46 vertebrate species. For the original EP calculations, we used species relationships from UCSC and divergence times from TimeTree web resource, and the resulting EP estimates were considered to be the ground truth. We found that the modified approaches produced reasonable EP estimates for HGMD disease missense variant and 1000 Genomes Project missense variant datasets. Our results showed that reliable estimates of EP can be obtained without a priori knowledge of the sequence phylogeny and divergence times. We also found that, in order to obtain robust EP estimates, it is important to assemble a dataset with many sequences, sampling from a diversity of species groups.

**Conclusion:** We conclude that the modified EP approach will be generally applicable for alignments and enable the detection of potentially neutral, deleterious, and adaptive alleles in populations.

## Background

The evolutionary probability (EP) method, introduced by Liu et al. [1], uses a Bayesian approach to produce a posterior probability of observation ranging from 0 to 1 for each possible allele at a site (e.g., each nucleotide for a DNA sequence, or each amino acid for a protein sequence). It requires a multiple species sequence alignment, phylogeny, and species divergence times. This method assumes no knowledge of the current state (i.e., allele) of the site in the species of interest, and relies solely on the observed configuration of alleles at the same site in other species in the sequence alignment. Low EP values indicate that an allele is unlikely to be common in a population of the focal species (evolutionarily forbidden alleles, eForb; EP < 0.05), whereas higher EP values indicate that an allele has been acceptable over the long-term history of species at the given position and may be more likely to be found (evolutionarily permissible alleles, ePerm; EP ≥ 0.05) [2]. Under the neutral theory framework, EP may serve as a null expectation for an allele’s frequency in a population, where alleles with high frequencies are expected to be ePerms and those with low frequencies are expected to be eForbs.

The EP approach has been applied to analyzing population polymorphisms in humans [1, 3], and the EP of alleles have been shown to correlate well with their population frequencies in the 1000 Genomes Project dataset for humans [1]. The EP approach is different from traditional methods (e.g., PAML [4] and HyPhy [5] software), because EP does not require to contrast patterns of synonymous and nonsynonymous changes. Also, the traditional methods do not use population frequency in designating adaptive changes. Thus, the EP approach complements other methods and provides site-by-site measurement of evolutionary estimates of neutrality of alternative alleles, based on multi-sequence alignments without requiring knowledge of synonymous changes, while incorporating population level information in making such determinations.

An analysis of a database of Mendelian disease associated missense variants (HGMD) showed that >90% of these variants are eForbs. Indeed, these disease-associated variants segregate with very low allele frequencies in humans. However, Patel et al. [3] previously reported more than 18,000 eForbs to be common in humans (allele frequency > 5%). They refer to them as candidate adaptive polymorphisms (CAPs), a collection which is likely enriched with truly adaptive alleles since it is comprised of eForbs with exceptionally high frequency. This CAPs catalog also contains a vast majority of known missense adaptive variants, which means that the EP approach is useful for forming hypotheses regarding natural selection at the molecular level.

The EP approach, however, has only been used for the above mentioned human datasets to date, even though it can be utilized for any species. This is partly because the application of the EP method to a multiple sequence alignment requires knowledge of the evolutionary relationship among sequences (phylogeny) and the divergence times for all the internal nodes in the phylogeny (timetree) [1]. For the analysis of human (and some other species, e.g., Fig. 1) proteins, such information is readily available from independent sources: for example, an evolutionary tree from the UCSC database and divergence times from the TimeTree resource [6, 7]. Such information is not as readily available for many other biological datasets, which discourages a more general use of the current EP method. Here, we present a modified EP approach in which the phylogeny and timetree are inferred from the sequence alignment and then the EP formulation of Liu et al. [1] is applied.

**Figure 1.**
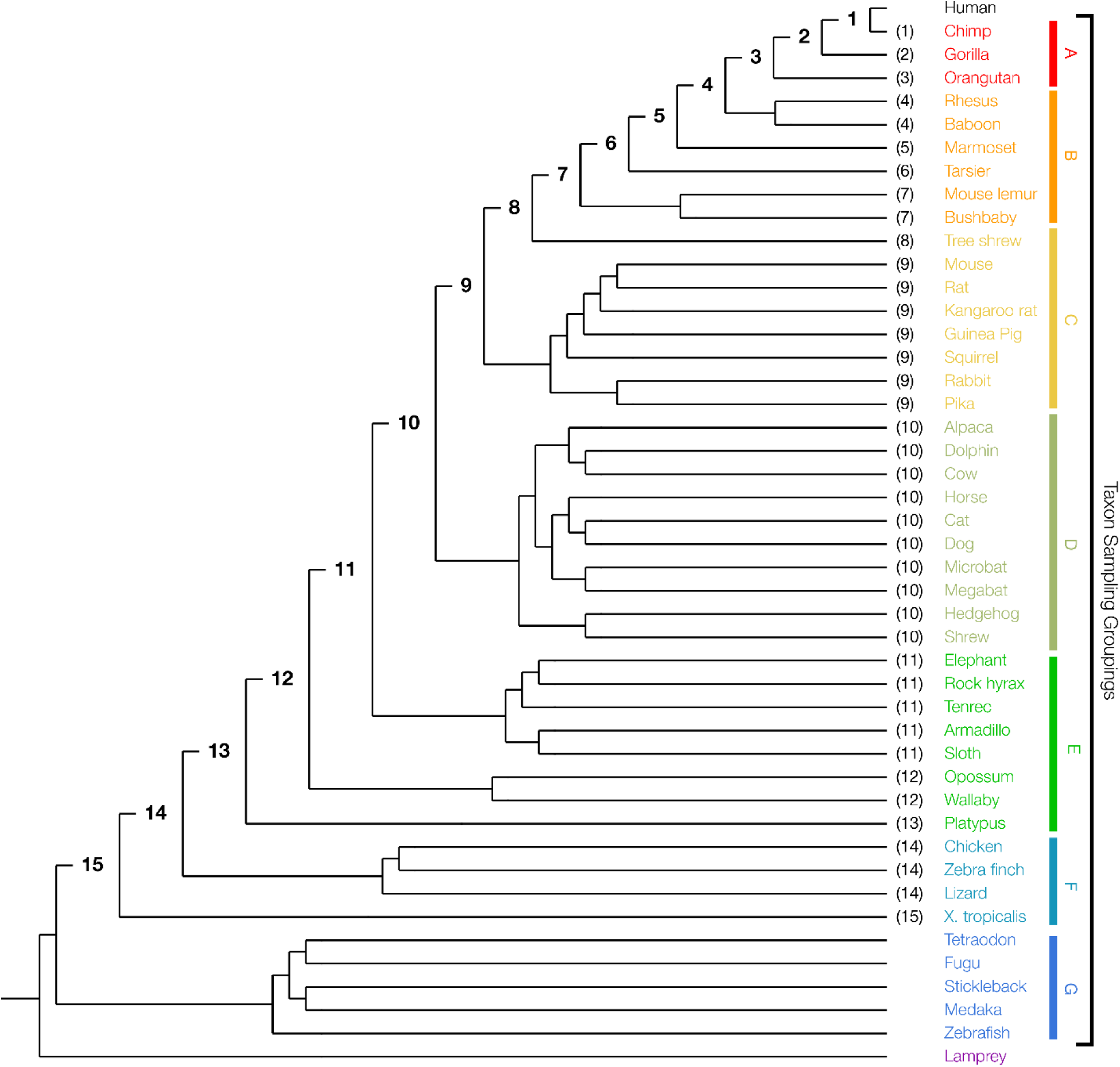
Phylogenetic relationships of 46 vertebrate species used for calculating evolutionary probabilities (EP). Nodes ancestral to the focal species, human, are labeled with numbers that correspond to pruning steps in EP calculation algorithm (see Methods). Numbers in parentheses next to the species label represent the step at which the taxon is pruned from the tree. Each of the seven main species groups used in the taxon density sampling are colorized (including the outgroup, lamprey) and labelled.

We evaluated the accuracy of the modified EP approach in discovering eForbs, ePerms, and CAPs by using the human protein variation data. This is because variation in the human exome has been the focus of genomics research for decades, and has a large, high-quality, record of annotations as well as polymorphism data. In the following, we first present the modified approach and then compare its performance with the original method. We show that useful estimates of EPs can be derived without a priori knowledge of phylogeny and known divergence times, as the phylogeny and times inferred from the sequence alignment serves as a good substitute and produces reliable inference of evolutionary permissibility. In order to examine the effect of sequence diversity in the multiple sequence alignment on this inference of evolutionary permissibility, we assessed the impact of taxon sampling on EP calculation and found that, as long as sufficient phylogenetic signal is present in the dataset, EP values produced by the modified EP approach are very similar to those from the original EP method. Therefore, the modified EP approach will be generally applicable for analyzing population variation in the context of multispecies and multigene family evolution.

## Materials and Methods

### EP methods

Evolutionary Probability captures neutral expectations for observing an allele by using a Bayesian analysis of long-term evolutionary history of the sequence. Using a multi-species alignment and phylogenetic relationships among the sequences, Liu et al.’s method [1] first estimates the posterior probability of observing any allele in sequence of interest by using the prior knowledge of the relationship among sequences and the sequences themselves. For example, EP can answer the question: “what is the probability of observing an alanine residue at amino acid position 42 in the *human* beta globin protein (HBB), given the multiple sequence alignment for *HBB* in 46 vertebrate species?” To answer such a question, Liu et al.’s method assumes that the actual residue at position 42 in the human sequence is unknown, and produces probabilities for all alleles possible at the site (20 amino acids).

Formally, EP of an allele at a sequence position in a given species in a tree is the weighted mean of a set of posterior probabilities {*PP*_0_, *PP*_1_, *PP*_2_, ⋯, *PP*_*n*_} calculated from the sequence alignment and species phylogeny. *PP*_0_ is the posterior probability of observing a specific allele at a specific position in the focal species where the full dataset is used. Here 0 indicates no sequences are excluded. *PP*_1_ is the posterior probability of the same allele at the same position after excluding the sister species or group closest to the focal species. The 1 indicates that the first closest group to the focal species was excluded. In the phylogenetic tree in **Figure 1**, this means that the chimpanzee lineage is excluded when computing *PP*_1_. This process is repeated for the residual phylogeny, which results in fewer species in progressive pruning steps. The pruning stops when the tree has only one outgroup and the focal species. The number of pruning steps (*n*) depends on the tree topology and the number of sequences in the tree. **Figure 1**, shows a total of 15 pruning steps for the 46 vertebrate species phylogeny, with humans as the focal species.

The weights of PPs used to calculate EP are the set of divergence times {*T*_0_, *T*_1_, *T*_2_, ⋯, *T*_*n*_}, where *T*_*i*_ for all *i* ≥ 0 is the divergence time between the focal species and the closest related taxon in the phylogeny used for calculating *PP*_*i*_. Then, using a standard weighted mean formulation:

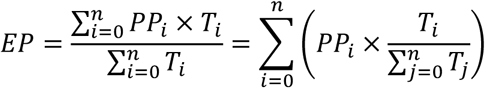

Therefore, the weights for posterior probabilities are normalized times, and are thus unitless.

The modified EP approach differs from the EP method of Liu et al. [1] in that the evolutionary relationships (phylogeny) of sequences in the given alignment and the divergence times among clades are both inferred from the sequence alignment itself. We suggest inferring such evolutionary relationships by using model-based methods, e.g., Maximum Likelihood under a suitable substitution model [8], which are known to be more accurate than the alternatives [9, 10]. In order to transform this phylogeny into a timetree, one may use a Bayesian method or a RelTime approach [11]. We selected RelTime, because its computational time requirements are orders of magnitude smaller [12]. Also, RelTime produces excellent relative times without requiring any calibration or other prior assumptions, as shown through extensive computer simulations [12, 13]. Additionally, the RelTime method has a strong theoretical foundation and produces results that are similar to those from Bayesian methods for empirical datasets [14–16]. These relative times can be directly used, because the weight function in the EP calculation effectively normalizes divergence times in the input, making relative and absolute times equivalent (see above). Thus, using either absolute times (as used in the Liu et al. application of EP) or relative divergence times (as used in this modification) in the calculations will produce identical results.

In the modified EP approach, however, we also used a modified weight for the EP calculations. Instead of the divergence time between the focal species and the closest related taxa, *T*_*i*_ is instead the evolutionary time span (ETS; see “Evolutionary Time Span” section) of the protein in tree at stage *i*. This approach is different from the Liu et al. implementation of EP, where later pruning steps were given higher weights because divergence times between the focal species and the closest related taxon increase in subsequent pruning steps. Here we decrease the relative contribution of later pruning steps because the an amino acid present in a distant taxon is less likely to be neutral than one observed in a closely-related taxon. The neutrality of an allele can be better estimated as information for more diverse and distant taxa are available at a site. As more taxa are included in a sample, a clearer picture of the results of natural selection can be gleaned.

We refer to the EP method where species relationships and divergence times used are known beforehand as the “original” EP method, and the EP method where species relationships and divergence times are both inferred as the “modified” EP approach.

### Data collection and analysis

We downloaded sequence alignments of 18,621 protein-coding gene orthologs in 46 vertebrate species from UCSC Genome Browser [17]. Where duplicate isoforms of the same protein were found, we selected the alignment with the longest sequence. We found that the sequences for 230 human proteins differed by >2% from RefSeq canonical sequences, so we excluded these from analyses. The remaining 18,391 sequence alignments were used to compute EP values for all tested approaches.

Missense variants used for evolutionary permissibility classification were acquired from the 1000 Genomes Project Phase III (1KG) dataset [18]. We found 543,220 sites at which a missense mutation occurs in at least one of the 2,504 individuals in the set of 18,391 proteins analyzed. For each protein, we computed amino acid EP values using MEGAX [19] under a Poisson model with a discrete Gamma distribution of rates (5 categories) that includes invariant sites (G+I). Other models could have been specified, but the estimates of EP were previously shown to be robust to the complexity of substitution model used [1]. For analyses where the phylogeny was presumed to be unknown, we first calculated maximum-likelihood trees in MEGAX using the same substitution models used in the EP calculation; branch lengths were discarded and only the topology was used.

Our human disease dataset consists of 50,422 disease associated missense variants retrieved from the Human Gene Mutation Database (HGMD) [20]. EP for each variant was calculated using both the original and modified EP methods described above.

### Evolutionary time span

A protein’s evolutionary time span (ETS) is the average of positional time spans (PTS) across all sites in a protein sequence alignment. PTS at a site is the total time along all branches in a tree for which a valid base (or residue, depending on whether nucleotide or protein sequence alignment) has existed in the evolutionary history of the site [21]. Alignment gaps and missing data in a multiple sequence alignment are not considered valid bases. To compute PTS for a site in a sequence alignment, the master timetree (used in the original EP calculation) is pruned such that only taxa that have a valid base at that site are retained. PTS is then simply the total time spanned by the resulting timetree (sum of times spanned by each branch) for that site. PTS will be a maximum for a site which has a valid base for all taxa in the master timetree.

Residue evolutionary time span (RTS) is the total time that a specific residue has been found in the evolutionary history of a site [22]. RTS is calculated by pruning the master timetree such that only taxa that possess only the specified residue are retained. RTS is the total time spanned by the resulting timetree (sum of times spanned by each branch) of a residue at a site. A residue that is not found in any sequence at a site has RTS of 0. RTS for all amino acids at a site will sum to the PTS for that site. A relative residue time span is often more informative than simple RTS, because it accounts for the PTS of a site and allows for comparison between sites with different PTS.

ETS can serve as a proxy for the amount of sequence information available; ETS that is close to the maximum indicates that there are few gaps in the sequence alignment, while ETS that is much lower than the maximum indicates a larger number of alignment gaps. PTS can convey similar information at the per-site level. Similarly, a small RTS means that the residue was found in a limited number of species and occupied that position for a limited amount of evolutionary time. In contrast, a large RTS means that the residue is commonly observed among species. Thus, time spans can be more informative to the properties of a sequence alignment as a relative value. So, here, we refer to all time span values as fractions of the maximum possible value of that measure (%ETS, %PTS, %RTS); i.e., %ETS is the proportion of a sequence alignment with no invalid bases covered by the ETS of the protein (ETS / maximum possible ETS), %PTS is the proportion of the time span covered by PTS for a site with valid bases for all species in the alignment (PTS / maximum possible PTS), and %RTS is the proportion of the PTS spanned by a specific allele (RTS / PTS).

### Tree distance

Branch-length distance [23] was used to quantify the error in inferred phylogenies, which were used in the modified EP analyses. The inferred tree was compared to the timetree used in the original EP method, but since the inferred tree produced relative time branch lengths, we first scaled the inferred tree such that its sum of branch lengths was equal to that of the original EP timetree. The branch-length distance, unlike simple symmetric differences or partition metrics, measures both differences in topology as well as branch length differences of the trees being compared. Such a measure is useful here because EP incorporates both species relationships (topology) and divergence times (branch lengths) into its calculations, so an ideal distance measure will capture differences in both of these properties.

### Taxon sampling

#### Sampling within clades

In our taxon “density sampling” experiments, the number of taxa included in each major clade of the 46 species vertebrate tree were varied (**Fig. 1**). We generated 100 replicate samples for one, two, three, and four taxa per clade (density) from seven clades (A-G, **Fig. 1**). Taxa were randomly sampled from these clades when generating replicate datasets, and humans were used as the focal species. For each analyzed clade density, the mean and standard error of EP were calculated for each residue, separately for original and modified approaches. Additionally, the mean ETS for all replicates was recorded for each clade density.

#### Sampling between clades

“Temporal sampling” iteratively increases the number of taxa distantly related to the focal species, human (**Fig. 1**). In each iteration, the next closest related taxon to the previous dataset is included. The first iteration requires a minimum of 3 taxa to analyze: human, chimpanzee, gorilla; the second iteration added orangutan, the fourth added rhesus monkey, until the final iteration contained all taxa including the lamprey.

### Receiver Operating Characteristic (ROC)

We calculated true eForb and false eForb classification rates under various eForb thresholds (EP value below which an allele is considered evolutionarily forbidden; 10 evenly spaced thresholds between EP<0.01 and EP<0.1) to determine the performance of the modified EP approach relative to the original EP method. For a given eForb threshold, we identified each eForb variant in the 1KG dataset based on EP values from the original EP method as the set of “condition positive”. 1KG variants that were not eForbs comprised the set of “condition negative” variants. For the same set of 1KG variants, we collected the set of eForbs identified across a variety of discrimination thresholds based on modified EP values as the set of “predicted condition positive” variants. Variants not predicted to be eForbs using modified EP values were the set of “predicted condition negative” variants. True(/false) eForb classification rates were calculated as the fraction of condition positive(/negative) variants that were correctly classified as eForbs(/not eForbs) when using the original EP values as the ground truth. ROC curves were generated for each of the eForb thresholds from 0.01 to 0.10, as described above.

## Results

The 1000 Genomes (1KG) dataset [18] contains sequence variation from 2,504 individuals. Among millions of variants present in this dataset, there are 543,220 missense variants that occur at non-zero population frequencies (**Fig. 2a**). We use this subset as our model and testing set. We consider the EP values obtained using the original EP method for these variants to be the ground truth, because the species phylogeny and divergence times used were not derived from any one protein alignment (as mentioned earlier). We computed EP values for 1,086,440 missense variants (major and minor alleles at missense sites; 2 × 543,200 missense sites) in the 1KG dataset using the original and modified EP methods. First, we examined the relationship between the EP value and population frequency of an allele. They are strongly correlated, similar to the pattern reported for the original EP method [1] (**Fig. 2b**). This is because of a strong agreement between the original EP values and modified EP values for human missense variants (*R*^2^ = 0.932). As mentioned earlier, the original EP method predicted evolutionarily forbidden (eForbs) alleles, which were important to diagnose disease-associated and detect putatively adaptive variants. So, we examined if eForbs identified using the modified EP approach produce results similar to the original EP method. Of the 1,086,440 missense variants in the 1KG dataset, 518,233 were classified as eForb by at least one of the EP methods (original or modified). The original EP method identified 494,821 eForbs, whereas the modified EP approach identified 508,065 eForbs (**Fig. 3a**). There was 93.5% agreement in that the original and modified EP methods both produced EP < 0.05 for a given method. Next, we evaluated if the modified EP approach performs as well as the original EP method in diagnosing 50,422 disease-associated missense variants found in HGMD. We found a 98.7% agreement, as the modified method designated 48,772 of HGMD variants to be eForbs, whereas the original method designated 48,657 of the HGMD variants to be eForbs (**Fig. 3b**). Overall, the low proportions of mismatched eForb designations suggest that the modified EP is a robust substitute for the original EP method, even when we use the topology and divergence times estimated from the sequence alignment.

**Figure 2.**
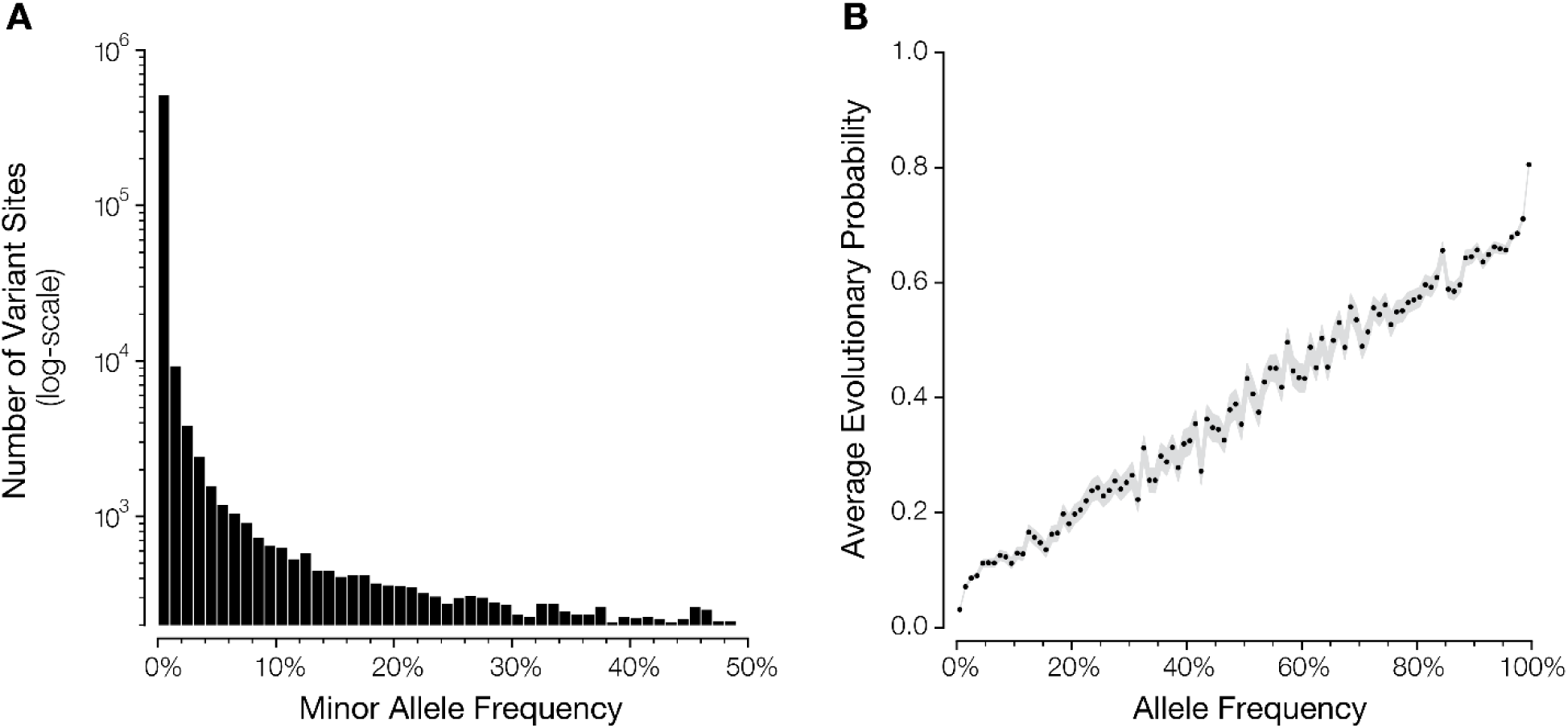
Population frequencies of missense sites found in 1000 Genomes Project Phase III dataset. (**a**) Distribution of minor allele frequency at positions containing missense variation. (**b**) The relationship between allele frequency (1% bins) and mean EP (modified method) of missense variants found in 1000 Genomes Phase III dataset. Gray area corresponds to standard error of the mean.

**Figure 3.**
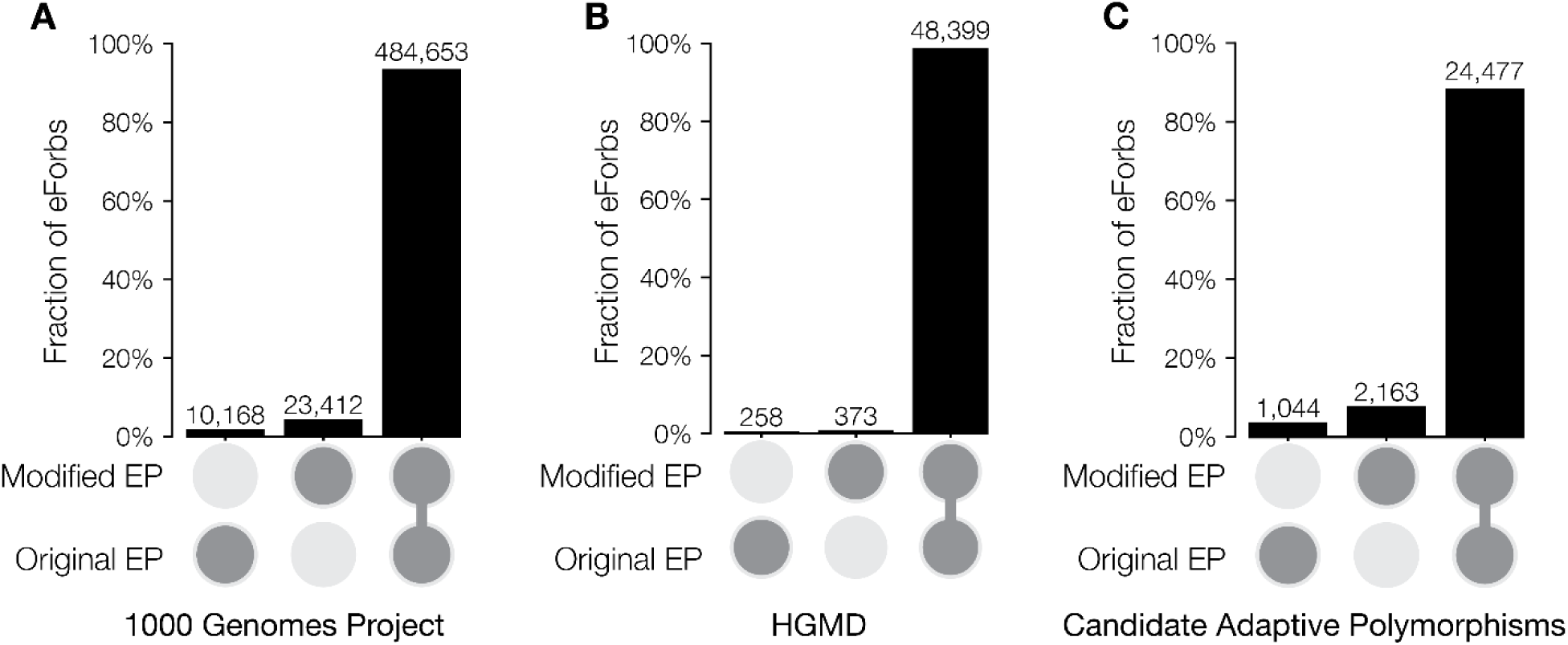
Designation of eForbs (EP < 0.05) using the original and modified EP methods. Agreement for classification of evolutionary forbidden alleles (eForbs) using the original and modified EP calculated methods for (**a**) all missense variants found in 1000 Genomes Project Phase III dataset, (**b**) human disease associated missense variants found in the HGMD disease variation dataset, and (**c**) high allele frequency (global AF > 5%) missense variants with EP < 0.05 (CAPs). Single darkened circles under a bar represent eForbs identified by the indicated method, and not the other. Connected darkened circles represent eForbs identified by both methods.

We also examined the eForb agreement between the two methods for variants found to occur at high allele frequencies (AF). eForbs segregating in the human populations at high AF (global AF ≥ 5%) are *candidate* adaptive polymorphisms (CAPs; [3]), because these variants are evolutionarily forbidden, yet segregating at unexpectedly high population frequencies, suggesting that some of them may have been positively selected. We again found high agreement (88.4%) between the two EP methods for identifying CAPs (high AF eForbs; **Fig. 3c**).

Furthermore, we similarly examined the handful of missense variants that are known to be adaptive in humans. As expected, given the strong concordance between the original and modified EP methods, the modified EP approach classified >95% (23/24) of these previously known adaptive missense alleles as eForbs (**Table 1**). One of these variants was not previously detected as eForb using the original EP method. Therefore, the new method is effective in identifying potentially adaptive variants.

**Table 1.** Known adaptive missense polymorphisms with their eForb status using both the Original and Modified EP methods. Table modified from Patel et al. [3].

| Protein | SNP Identifier | Phenotype | Is eForb? |  |
| --- | --- | --- | --- | --- |
|  |  |  | Original EP | Modified EP |
| ALMS1 | rs10193972 | insulin resistance | yes | yes |
|  | rs2056486 | insulin resistance | yes | yes |
|  | rs3813227 | insulin resistance | yes | yes |
|  | rs6546837 | insulin resistance | yes | yes |
|  | rs6546838 | insulin resistance | yes | yes |
|  | rs6546839 | insulin resistance | yes | yes |
|  | rs6724782 | insulin resistance | yes | yes |
| APOL1 | rs73885319 | Trypanosoma resistance | no | yes |
| DARC | rs12075 | malaria resistance | yes | yes |
| EDAR | rs3827760 | eccrine glands | yes | yes |
| G6PD | rs1050828 | malaria resistance | yes | yes |
|  | rs1050829 | malaria resistance | yes | yes |
| HBB | rs334 | malaria resistance | yes | yes |
| MC1R | rs1805007 | skin pigmentation | yes | yes |
|  | rs1805008 | skin pigmentation | yes | yes |
|  | rs885479 | skin pigmentation | yes | yes |
| SLC24A5 | rs1426654 | skin pigmentation | yes | yes |
| SLC45A2 | rs16891982 | skin pigmentation | yes | yes |
| TLR4 | rs4986790 | immune response | yes | yes |
|  | rs4986791 | pathogen recognition | yes | yes |
| TLR5 | rs5744174 | immune response | no | no |
| TRPV6 | rs4987657 | dietary calcium absorption | yes | yes |
|  | rs4987667 | dietary calcium absorption | yes | yes |
|  | rs4987682 | dietary calcium absorption | yes | yes |

### Causes of differences in eForb designation

While the two EP methods produce similar eForb designations, we investigated factors that may lead to some of the differences observed. Using the original EP method calculations, for which we had a known phylogeny and divergence time from independent sources, as the ground truth for designating eForbs, we scored alleles that did not receive an eForb designation by the modified approach. (We do not discuss the reverse scenario because the original method’s EP estimates are derived using more information [a priori phylogeny and times] than the modified approach.) For each protein, we computed the proportion of missense variants that were not classified (incorrectly so) to be eForbs by the modified EP approach (ΔeForb). ΔeForb for proteins range from 0 to ∼15% (**Fig. 4a**). That is, at most 15% of all alleles at polymorphic sites in a protein alignment were incorrectly classified as eForbs in any protein, although most proteins (82.2%) show ΔeForb < 5% (**Fig. 4a**). About half (52%) of proteins had no incorrectly classified eForb variants. A statistical test of gene ontology functional categories [24] did not find any biological process categories to be significantly over-represented, indicating that incorrect eForbs were not segregating in specific functional classes. Instead, ΔeForb was higher for proteins that evolved with faster evolutionary rates (**Fig. 4b**). We found that the sequence alignments of faster evolving proteins also tend to produce species trees that are increasingly different from the standard vertebrate tree used in the original EP calculation (**Fig. 4c and 4d**). Underlying this trend is the fact that even one substitution in a sequence can change the phylogeny topology relative to the standard vertebrate tree for highly conserved sequences, while sequence alignments for fast evolving proteins contain many more alignment gaps and missing data, and the proteins with the highest ΔeForb contained a large number of sites with alignment gaps (**Fig. 5a**). The impact of these alignment gaps is captured in the %ETS, which is a function of the prevalence of alignment gaps and missing data in an alignment that accounts for their evolutionary structure. The worst performing proteins had %ETS less than 50% (**Fig. 5a**). In other words, valid amino acid residues occupied positions for less than half of the total evolutionary time span possible in the vertebrate tree (2.84 byrs of 5.82 byrs) on average. We also observed a similar pattern for positional and residue ETS (%PTS and %RTS, respectively), namely that positions and residues that encompass larger timespans in the evolutionary tree produce the smallest ΔeForb (**Fig. 5b**, **Fig. 5c**).

**Figure 4.**
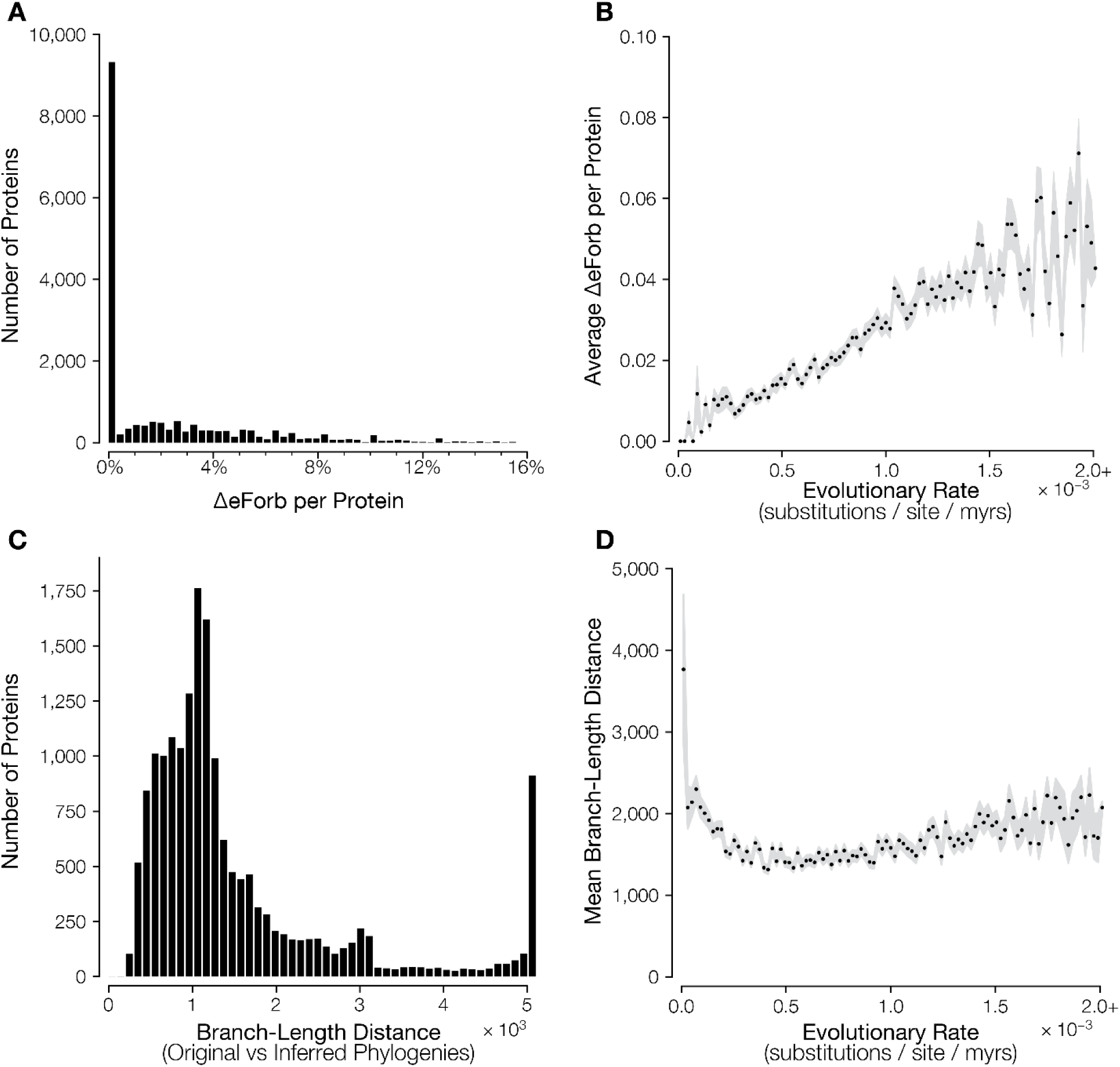
Relationship of protein evolutionary rate with eForbs classification error (ΔeForb). (**a**) Distribution of ΔeForb for 18,391 human proteins. (**b**) Proteins with higher evolutionary rates, on average, have higher ΔeForb. (**c**) The distribution of branch-length distances (tree difference) between the standard timetree and inferred RelTime trees. (**d**) Relationship between protein evolutionary rate and tree distance. For (**b**) and (**d**), the gray area corresponds to the standard error of the mean interval. Protein evolutionary rate is the ratio of sum of Maximum Likelihood estimates of branch lengths and the total evolutionary time in the tree of 46 species. Proteins with evolution rate >2×10^−3^ substitutions per site per million years were combined into one bin, shown as the rightmost points in panels (**b**) and (**d**).

**Figure 5.**
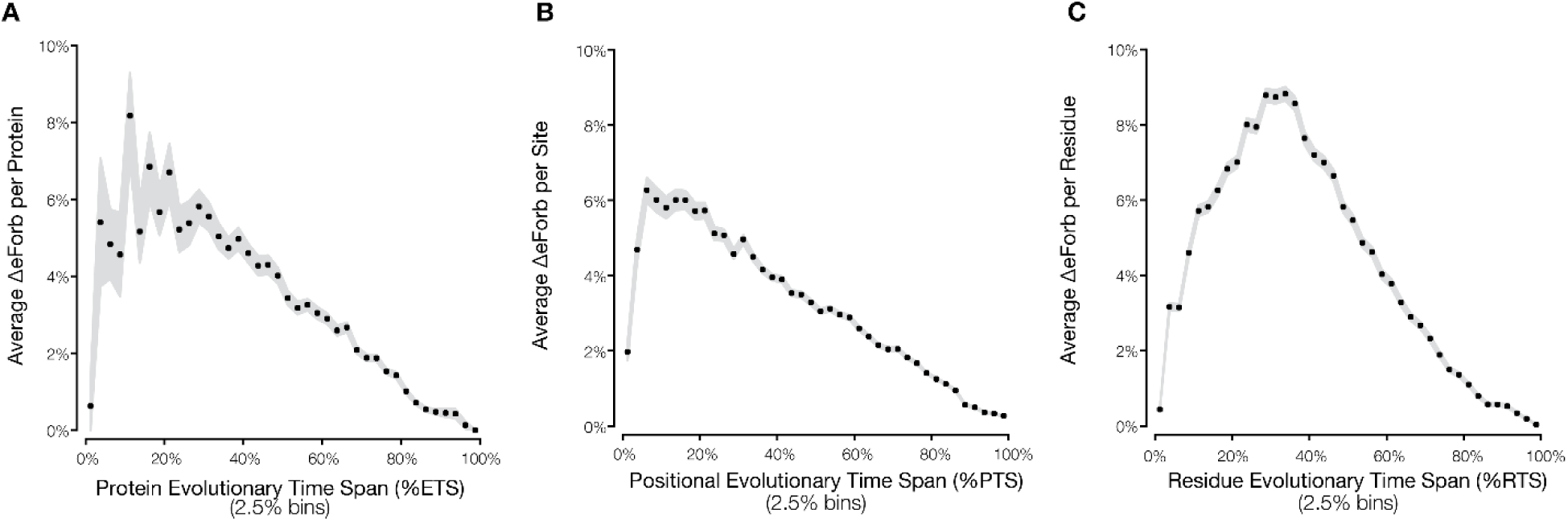
Error (ΔeForb) in designating eForbs by the modified EP method. Relationship of ΔeForb with (**a**) evolutionary time span (%ETS) of the whole protein, (**b**) positional time span (%PTS), and (**c**) residue time span (%RTS). For panels **a** and **b**, mean ΔeForb was estimated using values from all the positions in the specified time span bin. The maximum time span for %ETS and %PTS calculation is 5,819 million years (**Fig. 1**). Gray area represents the standard error of the mean.

While lower ΔeForb is correlated with higher %ETS, %PTS and %RTS, we find that ΔeForb can be low for positions with very low %ETS, %PTS and %RTS (**Fig. 5**). This is because amino acid residues with very low %RTS (e.g., < 15%) in the sequence alignment always produce low EP values since they are rarely observed among species. These EP estimates and thus eForb designations are not reliable whether we use the original or the modified method. Based on the trends seen in **Fig. 5**, it is best to trust eForb designations when the positions have relatively high %PTS. High %ETS alignments reduce error in EP estimated by the modified approach by producing better phylogenies than alignments with low %ETS. In fact, we found the phylogenetic error induced by low sequence coverage (time spans) to be the most important factor in ensuring concordance between the modified and the original EP approach. We investigated the effect of inferring only divergence times on EP values by using the correct species relationships (topology). Indeed, we found that EP values correlate strongly with the original EP values (*R*^2^ = 0.998; **Fig. 6b**), much better than the case where the phylogeny was inferred from the sequence alignment itself (**Fig. 6a**). Therefore, difficulty with phylogeny inference causes discordance between the original and modified methods, but the magnitude of the error is quite small in most cases.

**Figure 6.**
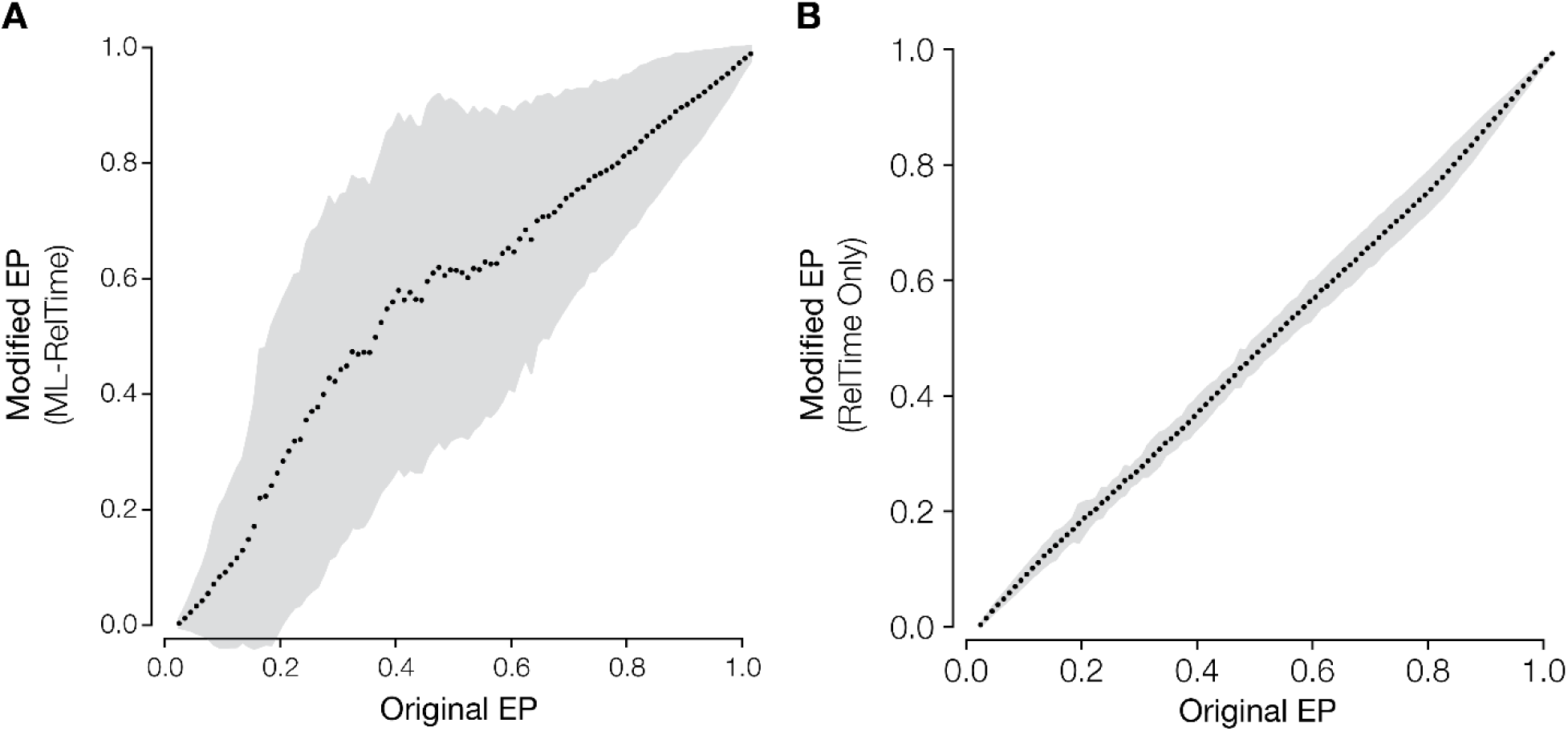
Evolutionary probability (EP) values for human missense variants using the standard and modified methods. The EP values on the x-axis are binned by 0.05 EP increments, with black points representing the mean EP of the (**a**) modified EP approach in which both species relationships and divergence times were estimated separately for each sequence alignment (ML-RelTime), and (**b**) modified EP approach in which only the divergence time was estimated and species relationships (**Fig. 1**) were assumed (RelTime Only). The gray areas represent the one standard deviation around the mean EP for the modified methods.

### Assembling a sufficient dataset

A researcher needs only a sequence alignment to apply the modified EP approach presented here. Inference of evolutionarily forbidden alleles requires a robust estimate of EP, which requires a sufficient sampling of sequences. The ultimate consideration for determining whether a dataset is sufficient is the total amount of evolutionary time spanned in the phylogenetic tree connecting the sequences (see “Evolutionary time span” in methods) because this will determine the number of mutations that have occurred or been “put to the test of natural selection” at a site. The more evolutionary time spanned in a tree, the more mutations will have occurred and been purged, or occurred and been allowed, at a given position in a sequence over evolutionary time. Alleles observed at a site will be the subset of mutations that were found to be acceptable. Thus allowing more time for mutations to have occurred at a site will increase confidence in alleles we consider evolutionarily forbidden; insufficient evolutionary time span will naturally lead to false eForb designations.

For many sets of species we can acquire evolutionary time spans from resources like TimeTree [6]. In such cases, researchers can determine whether sufficient evolutionary time has elapsed for a set of sequences by considering the per site mutation rate for the sequences of interest. For example, if we assume the DNA mutation for vertebrates to be the same as in mammals ∼2.2×10^−9^ per site per year [25], we can estimate the missense mutation rate per codon to be approximately ∼5×10^−9^ per year averaged over all possible trinucleotides. Given that a timetree of 46 vertebrate species spans ∼6 billion years, we expect each site to have experienced 30 missense mutations (= 6×10^9^ years × 5×10^−9^ missense mutations per year), which makes it highly likely that many different amino acids have been tested. Under these (idealized) conditions, if one or two residues dominate the position across vertebrates after ∼6 billion years, it is likely that most other alleles are unfavorable and, thus, can be inferred to be evolutionarily forbidden at that position. A tool to perform this estimation for various codon translation tables and custom mutation parameters is available online (see “Availability of data and materials”).

The evolutionary time span covered in a phylogeny can be increased either by sampling more taxa within clades already present in the sampled sequences (e.g., adding another primate to a set of mammalian sequences) or by sampling additional taxa from clades that are not present in the current sample of sequences (e.g., adding fish and bird sequences to a set of mammalian sequences). We expect the change in EP values per each additional sequence sampled to decrease, and thus, diminish improvement in identification of evolutionarily forbidden alleles. With this expectation, we investigated how the two approaches for expanding evolutionary time coverage impact inference of eForbs. Using the full species tree in the original EP method as the ground truth, we calculated EP using the modified method for a few select sites under various sub-samples of the full phylogeny. The temporal sampling scheme emulates the sampling of taxa from clades not already present in the phylogeny, while the density sampling scheme follows the approach of increasing sampling within clades already found in the phylogeny. Adding sequences under the former sampling scheme is expected to increase evolutionary time span faster than under the latter.

We focused on fast evolving sites because allelic EPs will be most impacted at these sites. EP estimation and eForb classification at completely and highly conserved sites is trivial, because only two EP values will be observed at such a site: ∼1 for the conserved residue, and ∼0 for all other unobserved (or rarely observed) residues. Fast evolving sites, however, will be especially sensitive to the sampled sequences and the specific configuration of alleles (i.e., which taxa possess each allele) among those sequences. For example, consider a fast evolving site, position 218 in human Poly (ADP-Ribose) Polymerase 9 protein, PARP9. It evolves 2.6 times faster than the average rate for the protein, and 5.6 times faster than the exome average. Under both sampling schemes, we found that certain alleles always maintain eForb status, regardless of the number of taxa sampled. These alleles are those that are never observed among the full vertebrate alignment, and are thus considered evolutionarily forbidden. There are others, however, that change from ePerm to eForb classification with increased evolutionary time span of the tree. For example, Glutamic acid (E) and Leucine (L) under a density sampling scheme (**Fig. 7**), and Glycine (G), Leucine (L) and Threonine (T) under temporal sampling scheme (**Fig. 8**). When the evolutionary time span is smaller, these residues are expected to be evolutionarily permissible, but their EP decreases as the evolutionary time span increases, which changes the classification ultimately to eForb. Slower evolving proteins will show similar patterns, but to a lesser degree.

**Figure 7.**
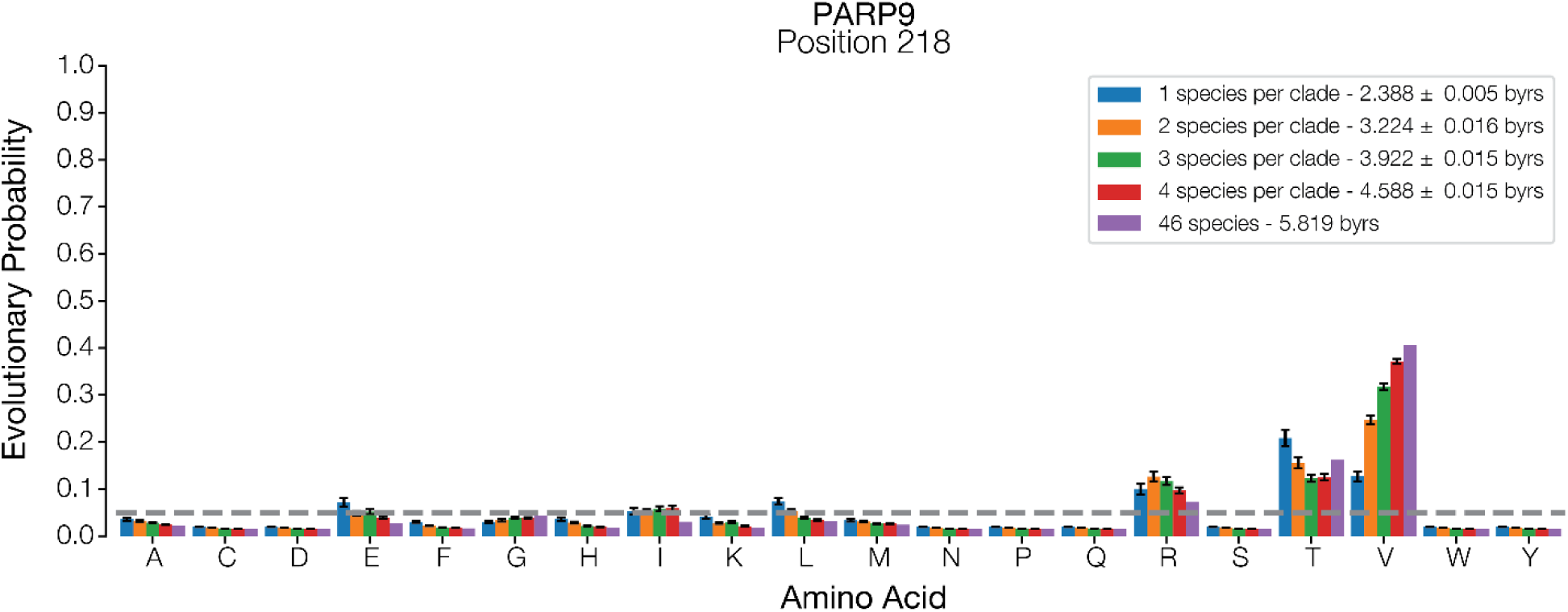
Effect of density sampling on EP value. Evolutionary probability (EP) values for each amino acid at position 218 in human Poly (ADP-Ribose) Polymerase 9 protein (PARP9) are shown for different taxa samples such that fewer or many species were included in the same set of clades. Dashed line marks EP = 0.05. The legend shows the mean (± standard error) evolutionary time spanned for all replicates.

**Figure 8.**
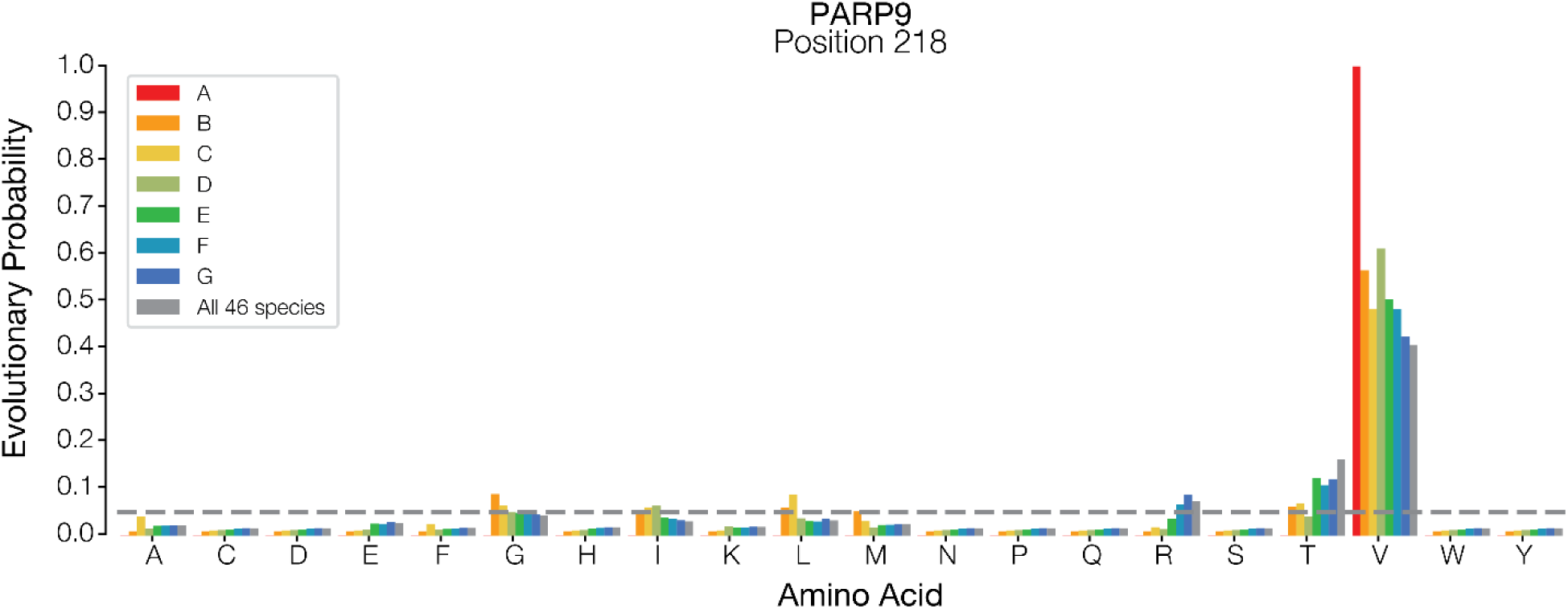
Effect of temporal sampling on EP estimates. Evolutionary probability (EP) values for each amino acid at position 218 in human Poly (ADP-Ribose) Polymerase 9 protein (PARP9) protein are shown for different taxon samples. Each bar represents an entire phylogenetic group that is sequentially sampled, such that all more closely related groups are included. Dashed line marks EP = 0.05. Colors and legend labels correspond to groups defined in **Figure 1**.

When too few distant taxa are sampled, we find that incorrect classification of eForbs is likely to occur, even when more evolutionary time is sampled than in a set of more distantly related taxa. For example, the Arginine (R) residue in our analysis is incorrectly classified as an eForb in the temporal sampling scheme even when 2.77 billion years of evolutionary history spanning all the mammals in the full tree is included in the EP calculations (**Fig. 7**). In contrast, sampling as few as seven total species that span 2.39 billion years of evolutionary history, one from each major clade in the analysis, correctly classified the Arginine residue to be evolutionary permissible (**Fig. 8**). Adding additional taxa to these clades does not change this classification. A similar result is observed for the Threonine (T) residue at this site.

While both sampling approaches show that incorrect eForb and ePerm classification can occur when too little evolutionary time is spanned by the sampled sequences, we do not find false eForbs when the evolutionary time is spread out over a variety of clades, instead of all compressed within a single clade; e.g., sampling 2 billion years of evolutionary time from a variety of vertebrates, instead of just from mammals, will lead to fewer incorrectly classified eForb residues.

## Discussion

In the presentation of the neutral theory, Kimura (1968) posited that the vast majority of substitutions observed among species were (nearly) neutral. From that, it follows that we can infer probabilities of observing various alleles under neutral evolution at a position by looking across species since the probability that an allele is neutral at a site increases as it is seen across more related species relative to those that are never observed. EP was proposed as a mathematical quantification of such relative probabilities [1], and happens to display characteristics that align with neutral theory expectations. First, detrimental alleles should not generally reach high AF in a population; in fact, we note a strong relationship between the EP of an allele and its AF in a population [3]. Specifically, low EP alleles have a low population AF, while high EP alleles have a high population AF. Second, a vast majority of known adaptive missense variants are found to have low EP. Similarly, human Mendelian-like diseases caused by missense variants are overwhelmingly due to low EP alleles (>98% of disease-associated alleles across all disease ontologies [2]). Together, these remarkable patterns suggest a straight-forward relationship between allelic neutrality and EP.

The ability to discriminate non-neutral (e.g., function-altering) alleles from those that have no impact on phenotype (neutral) is of high interest to researchers in diverse biological disciplines. EPs can be coupled with available polymorphism data to provide insight into detrimental and adaptive variants, as mentioned earlier. This approach is uniquely integrative, as other methods either focus on patterns among species only, or employ patterns of population variation to identify genes or genetic regions evolving adaptively [3]. While other methods have utilized the Empirical Bayes framework to infer probably sequences at various nodes in a phylogeny, e.g., ancestral sequence reconstruction [26, 27], the EP method is an advancement because it is explicitly designed to forecast contemporary sequences by uniquely incorporating the entire evolutionary history of a site. The weighting of the pruning steps in the modified EP provides a logical estimate of the permissibility of different alleles at a position, while remaining naïve to any phylogenetic signal in the contemporary sequence that would unduly influence inferences. Additionally, these methods are not robust to errors in phylogeny; that is, ancestral sequences are not useful if the relationship among species is not correct.

We have found the modified EP approach to perform well, i.e., estimation errors of phylogeny and divergence times have limited negative impact on EP estimates. This means that it can be widely applied, because unlike well-studied model organisms, where species relationships for related taxa are generally well resolved, phylogeny and times are known independently for only a small fraction of species. The modified EP approach was found to work well partly because the inferred species relationships from the sequence alignment themselves are not too different from the correct phylogeny. However, detecting eForbs reliably can be challenging when the sequence alignment contains a large number of insertion-deletions and missing data, which depletes the phylogenetic signal and evolutionary information. When a position contains a large number of alignment gaps and missing data, many residues would appear to be eForbs spuriously because of lack of sufficient information. This problem is more acute in the modified EP method, especially when the sequence alignment yields a phylogeny with a large number of errors. In such a situation, using a pre-determined phylogeny from another source, if possible, can help reduce error, as only divergence times will need to be inferred. Additionally, sites that are most phylogenetically informative [28] can be filtered prior to analysis to remove sites with low signal-to-noise ratio and help minimize errors in inference. Therefore, one needs to be circumspect when using EP estimates for positions with lots of missing data and alignment gaps, irrespective of the use of the standard or modified method.

In general, EP estimates can be improved by adding more sequences to the alignment. We explored two taxon sampling approaches to increase the total time spanned by a set of sequences. We found that sampling of additional species in clades not already present in phylogeny for sequences is more effective at increasing the evolutionary time span and decreasing error in eForb identification. While adding a taxon that is found in a species group already present in the tree will increase the total time span, it will result in a smaller total increase. So, adding new species groups is preferred over increasing the density of samples per group. In practice, we suggest adding as many sequences as possible, so denser and more diverse alignments are compiled for EP analysis.

Here, we have focused primarily on defining eForbs by assuming an EP threshold of 0.05. This threshold was found to be reasonable for humans given simulations of neutral sequence evolution in vertebrates [3]; i.e., a neutral allele was found to have EP < 0.05 at less than 1% of simulated sites. Given the strong relationship between EP values from the original and modified EP methods, the high success rates observed using the EP < 0.05 threshold is expected to hold regardless of the cutoff value. However, one might wish to use a more conservative or liberal approach and vary the EP threshold to designate eForbs. For the currently tested data, we compared eForb designations at different cut-off values by generating receiver operating characteristic (ROC) curves and calculating the area under the ROC curve (AUROC; see methods) using the standard EP method as the ground truth (**Fig. 9**). AUROC is very high (0.94) for EP < 0.05, and it remains high when we used a liberal cutoff of 0.10 (AUROC = 0.94) and when using a conservative cut-off 0.01 (AUC = 0.91). Thus, the EP approach reliably detects evolutionary forbidden alleles for a variety of evolutionary scenarios.

**Figure 9.**
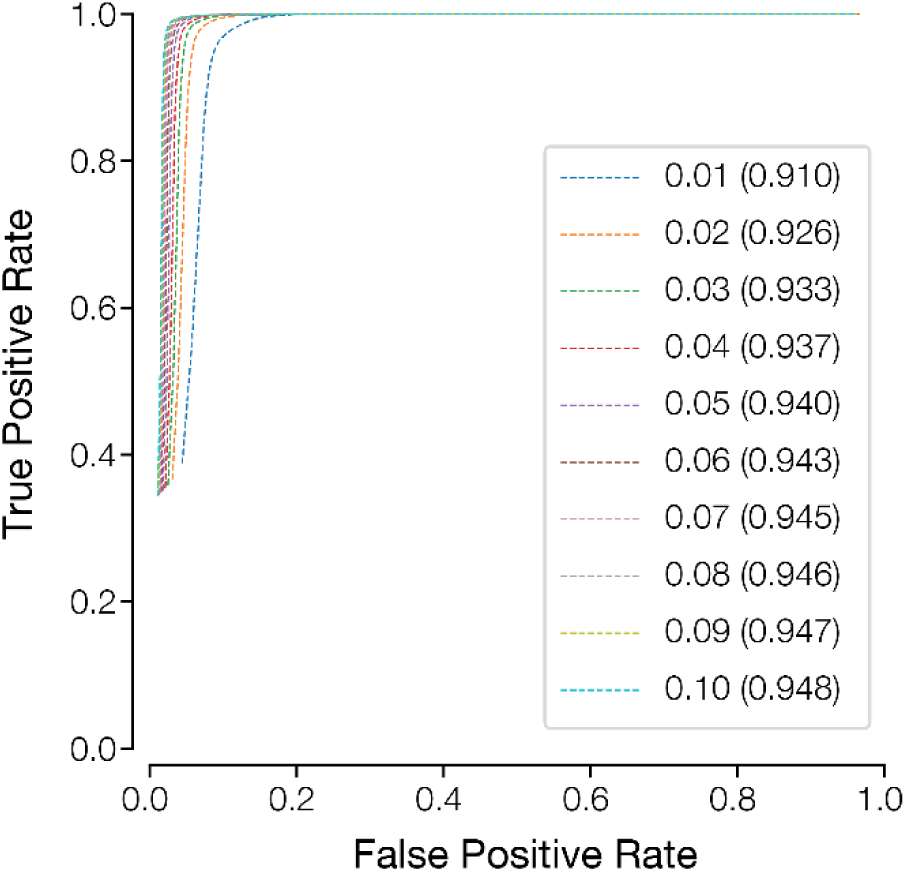
Receiver operating characteristic (ROC) curves showing the degree of misclassification caused by using EP threshold of 0.05 to designate eForbs, when the true EP thresholds for eForbs could be smaller (0.01) or higher (0.1). ROC curves are shown for classification of missense variants found in 1000 Genomes Project Phase 3 dataset using the modified EP approach with both species relationship and divergence times inferred from each sequence alignment. Area under ROC (AUROC) is shown in parentheses, which is similar for different thresholds.

## Conclusions

Evolutionary forbidden alleles can be predicted with high success even when the phylogeny and divergence times are estimated directly from the sequence alignment. It is, however, important that the species and genes included in the sequence alignment contain sufficient evolutionary information such that the expected number of mutations per position is as large as possible. This can be more easily accomplished by sampling sequences from distantly related species, as they add more evolutionary time span than the case where the taxon sampling is denser within each group. Of course, both approaches should be used whenever possible. With these alignments, one would be able to create catalogs of evolutionary permissible and forbidden variants for any gene or species, even when no polymorphism data exist.

## Declarations

### Ethics approval and consent to participate

Not applicable

### Consent for publication

Not applicable

### Availability of data and materials

Data sharing is not applicable to this article as no new datasets were generated or analyzed during the current study. Estimates of EP values are available for download from http://www.mypeg.info/epestimates. The modified EP approach is available for use in cross platform MEGA software (GUI and command line versions). A utility to estimate evolutionary time span required for robust EP estimation for sequence alignment using a variety of genetic codes and mutation rates is available at https://rpatel.github.io/ep-tools.

### Authors’ contributions

SK conceived the study, RP and SK designed analyses, RP conducted all the analyses, and RP and SK wrote the manuscript.

## Acknowledgements

We thank the following for their scientific feedback: Dr. Laura Scheinfeldt, Dr. Sayaka Miura, and Dr. Fernando Villanea.

